# Geometry-dependent instabilities in electrically excitable tissues

**DOI:** 10.1101/291617

**Authors:** Harold M. McNamara, Stephanie Dodson, Yi-Lin Huang, Evan W. Miller, Björn Sandstede, Adam E. Cohen

## Abstract

Little is known about how individual cells sense the macroscopic geometry of their tissue environment. Here we explore whether long-range electrical signaling can convey information on tissue geometry to influence electrical dynamics of individual cells. First, we studied an engineered electrically excitable cell line where all voltage-gated channels were well characterized. Cells grown in patterned islands of different shapes showed remarkably diverse firing patterns under otherwise identical conditions, including regular spiking, period-doubling alternans, and arrhythmic firing. A Hodgkin-Huxley numerical model quantitatively reproduced these effects, showing how the macroscopic geometry affected the single-cell electrophysiology via the influence of gap junction-mediated electrical coupling. Qualitatively similar geometry dependent dynamics were experimentally observed in human induced pluripotent stem cell (iPSC)-derived cardiomyocytes. The cardiac results urge caution in translating observations of arrhythmia *in vitro* to predictions *in vivo* where the tissue geometry is very different. We present simulation results and scaling arguments which explore how to extrapolate electrophysiological measurements between tissues with different geometries and different gap junction couplings.

## Introduction

Cells in multicellular organisms sense their location within tissues via diffusible molecules, contact interactions, and mechanical signals. Gap junction-mediated electrical signals can also, in principle, provide long-range positional cues ^1^, though mechanistic details have been difficult to determine due to the simultaneous presence of, and interactions between, all of the above signaling modalities in physiological tissue. Furthermore, until recently, technical limitations prevented tissue-scale mapping of membrane voltage: point-wise measurements with patch pipettes were slow and laborious, and voltage-sensitive dyes lacked sensitivity and suffered from phototoxicity.

The electrophysiological properties of many isolated cells have been probed in great detail via patch clamp electrophysiology.^2^ In tissues, cells form electrical connections with their neighbors via gap junction channels. One can then ask whether this coupling is a minor perturbation on the individual cells, or whether it fundamentally changes the dynamics. In condensed matter physics, the properties of a bulk solid can differ dramatically from those of its constituent atoms. Similarly, the emergent electrical properties of bulk tissue might differ dramatically from those of individual cells.

One aspect of positional sensing is the detection of boundaries. It has been well established that boundaries can influence paracrine signaling pathways^3^, but it remains an open question to what extent tissue geometry and topology influence electrical signaling. In the heart, structural defects can act as nuclei for arrhythmias ^4^, but interpreting these effects is difficult due to the multiple interacting factors that govern dynamics, including electrical coupling between myocytes ^5-7^, mechano-electrical feedbacks ^8, 9^, and differences in cell-autonomous properties of the individual myocytes between regions of the heart ^10-12^. Subtle shifts in any of these parameters can cause discontinuous changes in dynamics, e.g. from a stable beat to a possibly fatal arrhythmia.

It is important to understand how long-range electrical signaling can convey positional cues generally, and how these signals govern cardiac stability specifically. Early models of cardiac dynamics established a stability criterion based solely on the cell-autonomous relation of spike width to beat rate.^13^ Recent theoretical work showed that conduction could dramatically alter the stability conditions.^14, 15^ However, the wide diversity of cardiac models, combined with uncertainty in model parameters, presents a challenge for comparison to experiments.^15, 16^ Only a few experiments have explicitly probed the roles of intercellular coupling in cardiac dynamics.^17, 18^ In complex cells such as cardiomyocytes, one typically cannot vary one parameter without affecting many others. For instance, growing hiPSC-CM on different size islands affects their patterns of gene expression ^19^, which in turn can affect electrophysiology.

Uncertainties regarding the role of geometry in cardiac stability have an important practical implication: it has been widely claimed that if human induced pluripotent stem cell (iPSC)-derived cardiomyocytes (hiPSC-CM) can be made to show mature patterns of ion channel expression,^20-23^ then *in vitro* cultures will be a useful substrate for studying arrhythmias.^24-27^ However, this claim might need to be reconsidered if one finds that there are fundamental geometry-driven differences in stability between cultured cells and intact tissue, even when all voltage-dependent conductances are identical.

To explore the role of geometry under controlled conditions, we engineered a synthetic excitable tissue where all elements were well understood. This synthetic approach has the further merit of being amenable to rigorous quantitative modeling. We previously introduced Optopatch Spiking Human Embryonic Kidney (OS-HEK) cells^28^ as an engineered excitable cell type with an all-optical electrophysiological interface. The cells expressed just two voltage-dependent channels, the voltage-gated cardiac sodium channel, Na_V_ 1.5, and the inward rectifier potassium channel, K_ir_2.1. Expression of a channelrhodopsin permitted optogenetic stimulation, and expression of a far-red voltage-sensitive protein (QuasAr2^29^) or dye (BeRST1^30^) reported electrical dynamics. When optogenetically stimulated, these cells produced single action potentials. When grown into a confluent syncytium, endogenous gap junctions coupled cells to their neighbors, supporting bulk propagation of electrical waves.

While the OS-HEK cells demonstrated many interesting attributes of excitable tissues (including wave conduction, curvature-dependent wavefront velocity, and re-entrant spiral waves^28^), they did not show complex dynamical bifurcations (‘arrhythmias’) at fast pacing rates. Irregular and chaotic dynamics have not previously been observed in a synthetic bioelectrical system, suggesting that a necessary ingredient was missing.

Here we describe isradipine-OS-HEK (iOS-HEK) cells, a synthetic bioelectric system which shows dynamical transitions between regimes of regular pacing, complex but repeating patterns, and pseudo-chaotic (non-repeating) dynamics as the drive frequency is changed. We explore in detail how these stability regimes are influenced by the macroscopic tissue geometry. Remarkably, we found that the transitions to complex and irregular patterns depended sensitively on the culture geometry. At a single pacing frequency, we simultaneously observed regular rhythms, alternating patterns, chaos, or depolarization block in islands that were identical in all respects except for their geometry. A biophysically detailed Hodgkin Huxley-style model captured these geometric effects. The iOS-HEK cells further showed second-degree conduction block in regions of high wavefront curvature, demonstrating sensitivity to two-dimensional geometric features. Finally, we show that similar geometry-dependent transitions occur in cultured human iPSC-derived cardiomyocytes.

Together our findings show that macroscopic tissue geometry is a fundamental determinant of bioelectrical dynamics, and not just a perturbation on the cell-autonomous behavior. We discuss which parameters are sensitive or insensitive to tissue geometry, and propose scaling relations that can be used to extrapolate across tissue geometries and intercellular coupling strengths.

## Results

### iOS-HEK cells show alternans and arrhythmias

Our previous studies of OS-HEK cells did not show frequency-dependent stability transitions. We hypothesized that this was due to the absence of any slowly recovering conductances that could provide memory effects between excitations. Upon repolarization after a spike, OS-HEK cell ion channels recovered fully within 4 ms, so each beat was independent of its predecessors. The sodium channel blocker isradipine shows state-dependent block of Na_V_1.7, with a recovery time of 200 ms at –100 mV.^31^ We reasoned that in the presence of isradipine, the Na_V_1.5 channels in OS-HEK cells would show a similar slow recovery after a beat, introducing the possibility of complex temporal dynamics.

In a confluent monolayer of OS-HEK cells, we observed regular spiking when the cells were optogenetically paced at 4 Hz. Addition of isradipine (10 μM) converted the spiking to an alternating rhythm between large and small spikes (i.e., ‘alternans’, Fig. 1b), consistent with a previous report.^31^ We refer to the OS-HEK cells with 10 μM isradipine as iOS-HEK cells.

**Figure 1.**
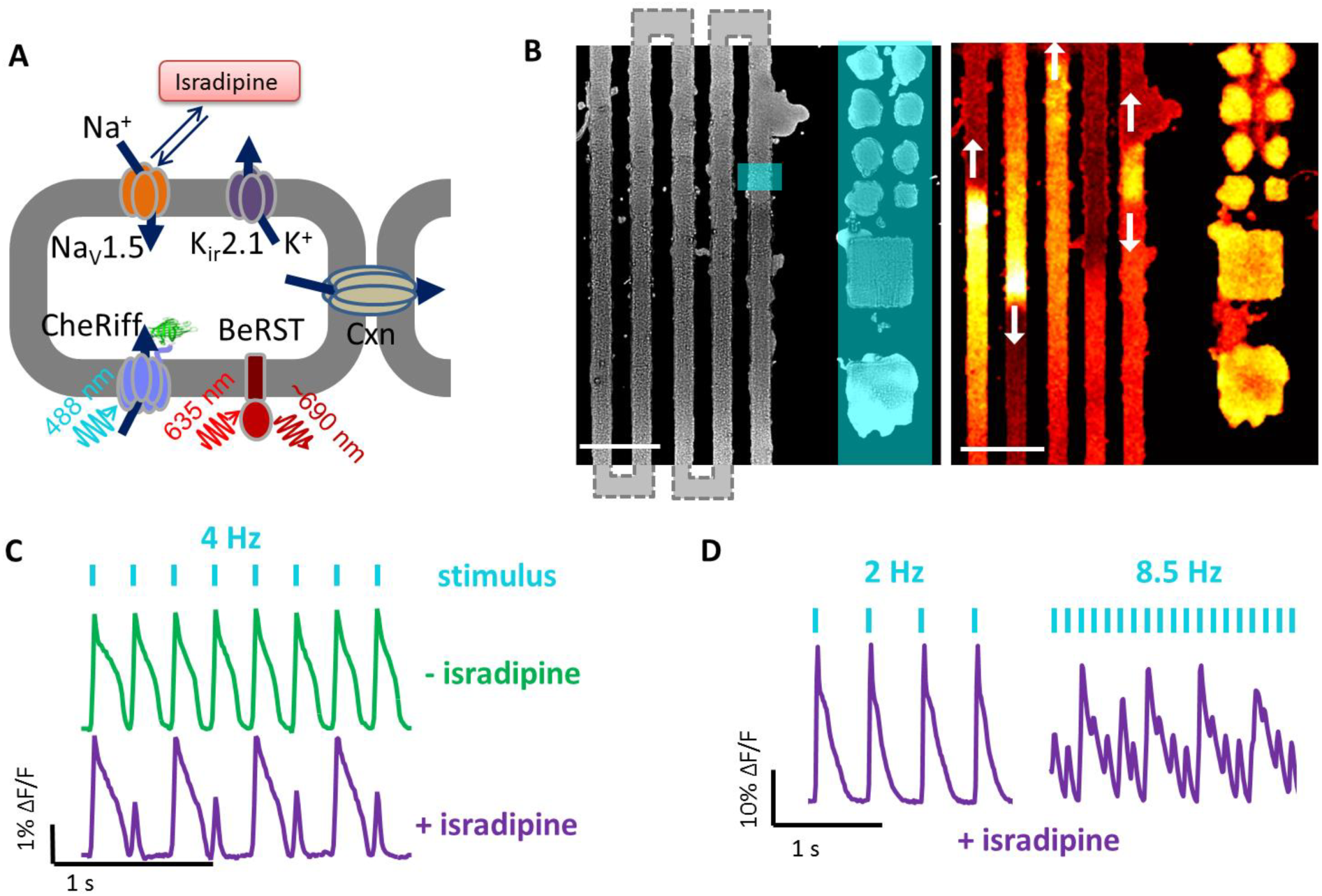
Isradipine renders Optopatch-Spiking HEK cells susceptible to arrhythmia. A) Molecular components of the synthetic excitable tissue. The voltage-gated sodium channel Na_V_1.5 and inward-rectifying potassium channel K_ir_2.1 are together sufficient to produce action potentials in response to a depolarizing stimulus. CheRiff is a blue-shifted channelrhodopsin which depolarizes the cells in response to blue light, and BeRST1 is a dye which reports voltage through changes in red fluorescence. Connexin channels (Cxn) introduce electrical coupling between neighboring cells. B) Left: microcontact printing defines patterns of cell growth. The vertical stripes connect in a serpentine pattern outside the field of view. Blue overlay shows regions of optogenetic stimulation. Zero-dimensional “islands” are tested alongside one-dimensional “tracks”. Right: single frame from a movie of BeRST1 fluorescence showing traveling waves in the track region. See **Supplementary Movie 1**. Scale bars 500 μm. C) Fluorescence recordings of membrane voltage showing that isradipine (10 μM) induces alternans in OS-HEK cells paced at 4 Hz. D) At low pace frequency isradipine OS-HEK (iOS-HEK) cells beat periodically in synchrony with the pacing, but at high pace frequency the cells produce an irregular rhythm.

Using microcontact printing,^32^ we patterned cell-adhesive fibronectin features onto cytophobic polyacrylamide surfaces. Features comprised square islands of linear size 100 μm, 200 μm and 500 μm, as well as serpentine tracks of width 100, 200, 500 μm and edge length of 1 mm and 5 mm. Overall track lengths were as long as 7 cm. Cell growth followed the printed patterns (Fig. 1c). We used a digital micromirror device (DMD) to target blue light stimulation to specific regions of the sample. In a change from previous studies on OS-HEK cells, we used the far red dye BeRST1 ^30^ to report membrane voltage (Fig. 1a). This dye had superior brightness to the protein-based reporter, QuasAr2, and showed otherwise similar response. We recorded the voltage dynamics using a custom ultrawide-field ‘Firefly’ microscope.^33^

Localized stimulation of the serpentine tracks induced propagating electrical waves (**Supplementary Movie 1**). Optogenetically induced waves propagated with a typical conduction velocity of 3.3 cm/s, had an AP width at 50% repolarization of 110 ms, and thus had a depolarized action potential length, *λ*, of *λ* = 3.6 mm. The fluorescence in the paced region of the track showed complex beat rate-dependent dynamics. At 2 Hz pacing, the cells spiked regularly and in phase with the drive (Fig. 1d, left). At 8.5 Hz pacing, the cells showed a complex and irregular fluorescence pattern, indicative of arrhythmia (Fig. 1d, right).

### Geometry dependent instabilities in iOS-HEK cells

We characterized in detail the dependence of the electrical dynamics on local geometry. Cells were grown either in small square ‘islands’ or on adjacent linear ‘tracks’, and paced simultaneously in both geometries at frequencies between 2 and 11 Hz. The islands were paced with spatially homogeneous illumination, so cells across each island spiked synchronously and gap junction-mediated conduction did not contribute to the dynamics. We refer to these as zero-dimensional (0D) dynamics.

Tracks were stimulated in small regions (200 μm wide, 100 μm long) to induce 1D propagating waves. We observed the response in both the directly stimulated region (near field) and in the conductively stimulated region (far field; Fig. 2a). Electrical waveforms stabilized to their far-field dynamics within a distance d ≈ 0.05 *λ* from the stimulus (d = 180 μm in our experiments), so we characterized the far-field dynamics at a distance 750 μm from the stimulus. Island and track features were intermixed within each dish, were seeded with the same stock of OS-HEK cells, and were measured simultaneously. Thus any differences in dynamics between regions could be ascribed to the island geometry.

**Figure 2.**
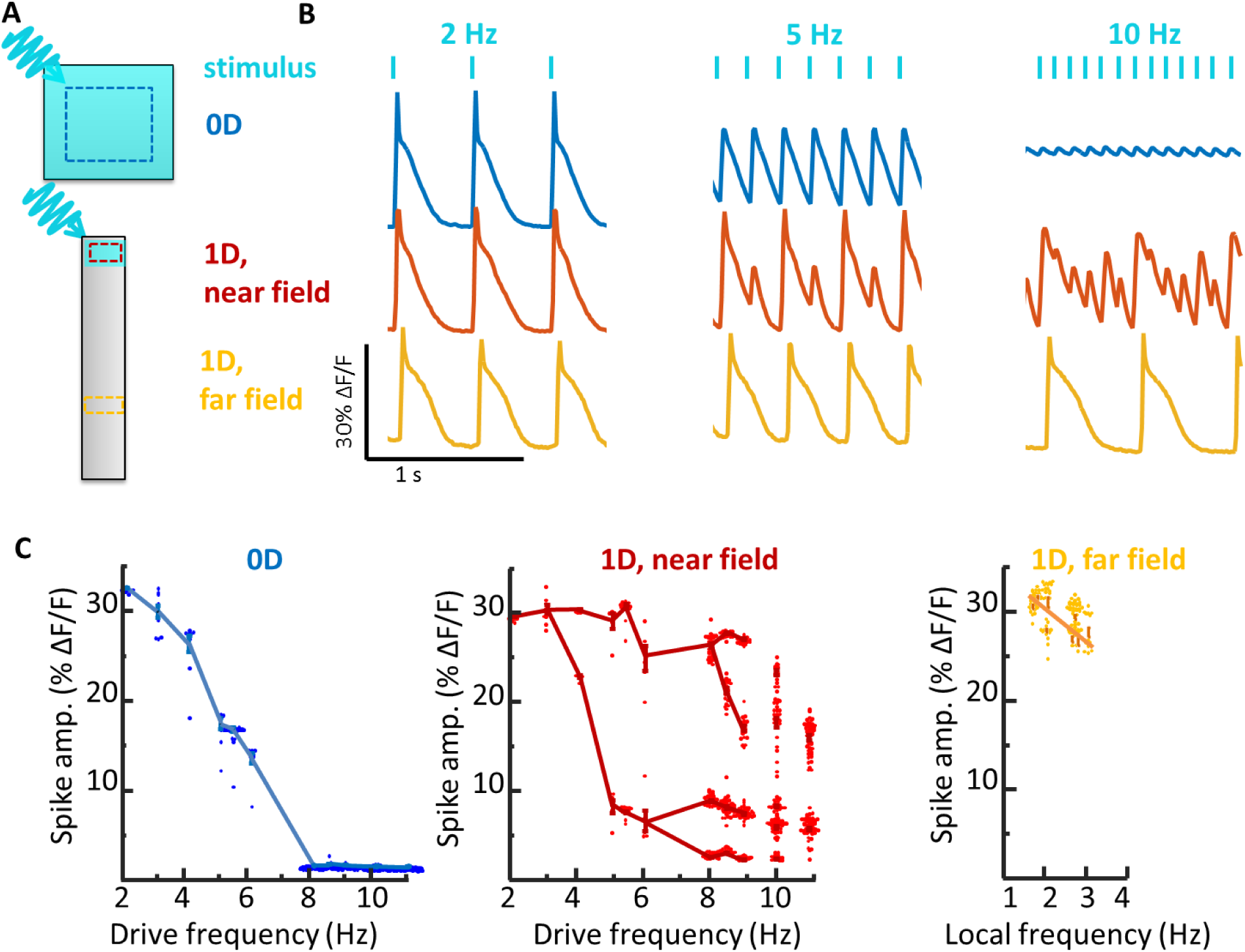
Stability of spiking in iOS-HEKs cells depends on sample geometry and location of pacing. A) Cell culture geometries, with locations of optogenetic stimulus shown in blue and regions of fluorescence voltage imaging shown with dotted lines. See **Supplementary Movie 2**. B) Simultaneously recorded electrical dynamics in 0D, 1D near field, and 1D far field. At pacing frequencies of 5 and 10 Hz the cells showed different dynamics in the three regions. C) Geometry-dependent spike amplitude as a function of beat frequency. The plots for 0D and 1D near field show spike amplitude as a function of pacing frequency. The plot for 1D far field shows spike amplitude as a function of local frequency, which can be a sub-harmonic of the pacing frequency.

At low pace frequencies (≤ 3 Hz), all three regions spiked with a stable rhythm in synchrony with the pacing (Fig. 2b). To our surprise, at higher pace frequencies we observed dramatically different dynamics in the three regions (**Supplementary Movie 2**). The 0D islands always produced regular electrical oscillations at the pace frequency. As the pace frequency increased, the amplitude of these oscillations diminished, up to 8 Hz, beyond which the responses were undetectable (Fig. 2).

The 1D near-field response developed a 1:1 alternans pattern at frequencies > 3 Hz. Additional bifurcations arose at 6 Hz and 8 Hz. At 10 Hz the dynamics no longer showed any repeating pattern, suggesting a transition to chaos (Fig. 2).

Remarkably, the 1D far-field showed no transitions to alternans or chaos (Fig. 2b). All spikes in the far-field had nearly equal amplitude and waveform. At pace frequencies between 3 and 6 Hz, only every other spike propagated to the far field, i.e. the local beat frequency was half the pace frequency. At higher pace frequencies, a smaller portion of spikes reached the far field. These spikes had irregular timing in the near-field but conducted at velocities which gradually evened out timing variations, such that the spiking appeared regular in the far-field, always at a frequency < 3 Hz. The 1D track acted as a filter which converted high-frequency arrhythmic spiking in the near-field into lower frequency rhythmic spiking in the far field. Together these experiments gave the unanticipated result that under regular pacing, iOS-HEK cells showed irregular dynamics only in the 1D near-field, not in the 0D or 1D far-field regions.

### A Hodgkin-Huxley model captures iOS-HEK dynamics

We developed a conductance-based Hodgkin Huxley-type model to simulate the dynamics of iOS-HEK cells. The properties of Na_V_1.5, K_ir_2.1, and channelrhodopsin CheRiff are all well known, so we were able to constrain the model with a small number of free parameters (Fig. 3a). The sodium channel model comprised Hodgkin-Huxley activation and inactivation gates *m* and *h* with dynamics taken from the literature.^34, 35^ To capture the use-dependent block by isradipine, we introduced an additional slowly activating and slowly recovering gate, *j* (Methods). The K_ir_2.1 conductance was modeled as an instantaneous function of voltage, inferred from the shape of the action potential repolarization (Methods). The channelrhodopsin was modeled as a linear conductance with a 0 mV reversal potential, modulated in space and time by the blue light illumination. A diffusive term captured the nearest-neighbor gap junction coupling. The governing equation is:

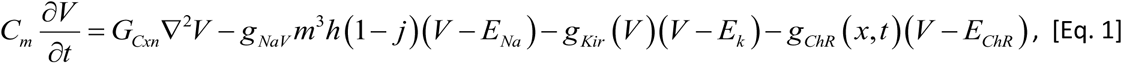

where *C_m_* is the membrane capacitance of a cell, and *G_Cxn_* = *g_Cxn_* ×*l^2^*, where *g_Cxn_* is the gap junction conductance between cells, and *l* is the linear dimension of a cell. The electrical diffusion coefficient is given by *G_Cxn_ / C_m_*. To simulate 0D dynamics, *g_Cxn_* was set to zero. Eq. 1 represents a continuum model, which treats the domain as homogeneous tissue. In simulations, the cell discreteness is recovered by setting the spatial discretization equal to cell length.

**Figure 3:**
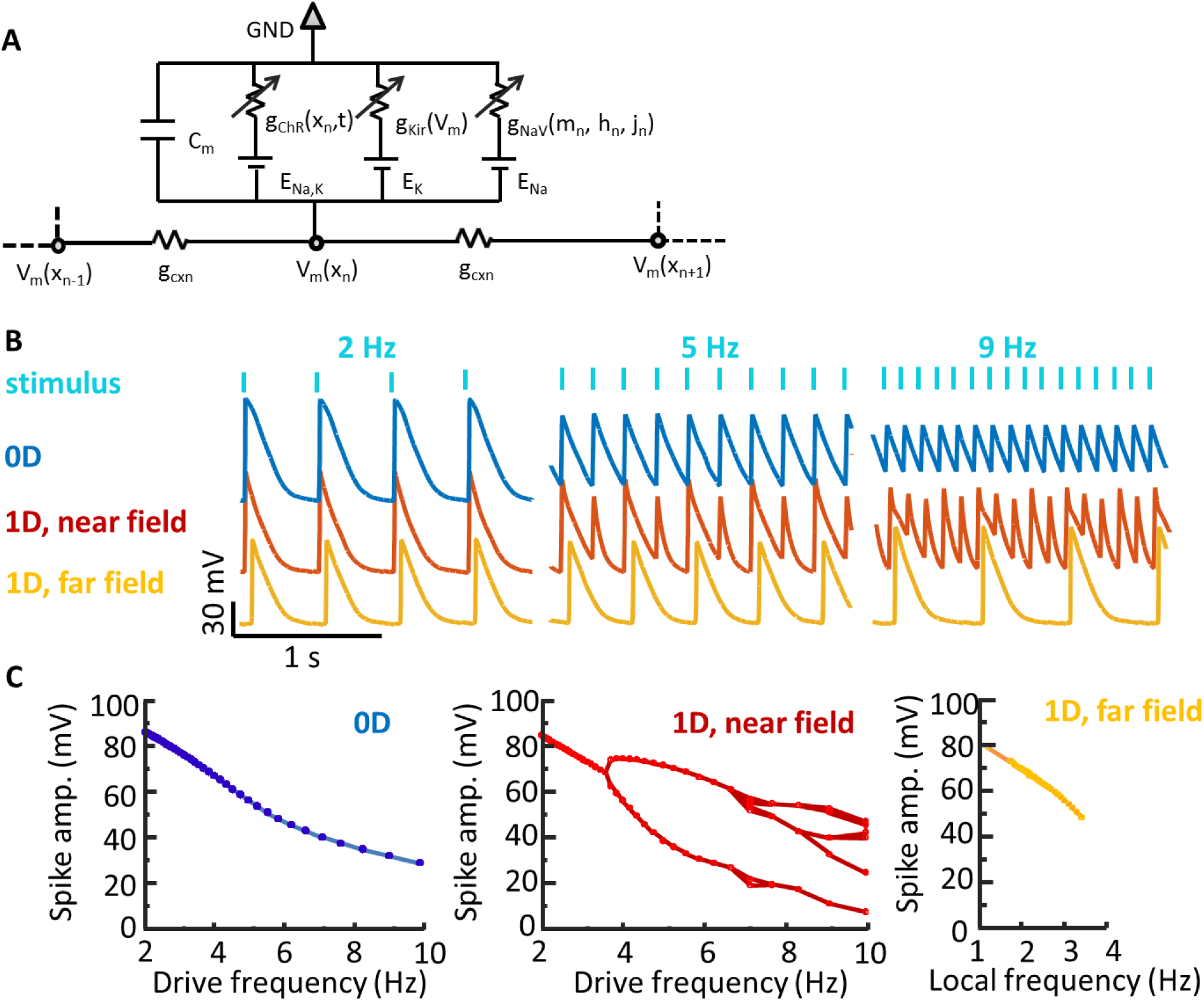
A Hodgkin-Huxley model recapitulates the effect of geometry on spike dynamics. A) Model schematic. Each cell has a channelrhodopsin (ChR), an inward-rectifier potassium channel (K_ir_ 2.1) and a voltage-gated sodium channel (Na_V_ 1.5). The Na_V_ is further gated by a state-dependent isradipine block. Neighboring cells are coupled via gap junctions. B) Simulated action potential waveforms in different geometries and pace frequencies (analogous to data in Fig. 2b). C) Simulated spike amplitude as a function of frequency for three geometries (analogous to data in Fig. 2c).

The unknown parameters (*C_m_, g_Cxn_, g_cKir_, g_ChR_*) were determined by fitting to observed fluorescence waveforms, conduction velocities, and patch clamp measurements (Methods). The free parameters in the model enter in the dynamics of the isradipine variable, *j,* which was assumed to bind sodium channels in their open state with rate *α,* and unbind with rate *μ*:

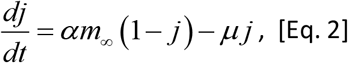

where *m*_∞_ is the steady state value of the *m* gate. The kinetic parameters in Eq. 2 were chosen to fit the experimentally observed dynamical transitions.

Action potential waveforms were simulated in a 0D geometry and in a linear 1D track comprising 2000 cells, with pacing delivered to 40 cells on one end. The near-field response was monitored in the paced zone, and the far-field response was monitored at a distance of 3 mm from the stimulus.

Simulated electrical waveforms (Fig. 3b) captured the main geometry and frequency-dependent features of the data. Specifically, the simulations in 0D showed a smooth and monotonic decrease in spike amplitude with increasing pace frequency. Simulations in the 1D near-field showed a series of frequency-dependent bifurcations that became increasingly irregular at high pace frequency. In the 1D far-field, these bifurcations were suppressed: spikes that propagated to the far-field had full amplitude, regular spacing, and were within the narrow band of frequencies that supported far-field propagation. These simulations confirm that cells with identical conductances, paced at the same frequency, can show widely divergent behavior depending upon the geometry of the surrounding excitable tissue.

### Mapping the transition between the near field and far field response

We next investigated the transition from near-field to far-field behavior. How does alternans, or even chaos, in the near-field lead to regular spiking in the far-field? We mapped the fluorescence dynamics of 1D tracks as a function of distance from the stimulated zone, across a range of stimulus frequencies (Fig. 4a,b). There was a clear bifurcation in the ability of spikes to propagate into the far-field. Near-field spikes with amplitude lower than a critical threshold decayed as a function of distance, while spikes above this threshold grew to become propagating far-field spikes. Spatially resolved simulations yielded similar results (Fig. 4d). The simulations showed that the conduction velocity of each far-field spike decreased as it approached the preceding spike such that irregularities spike spacing gradually evened out.

**Figure 4.**
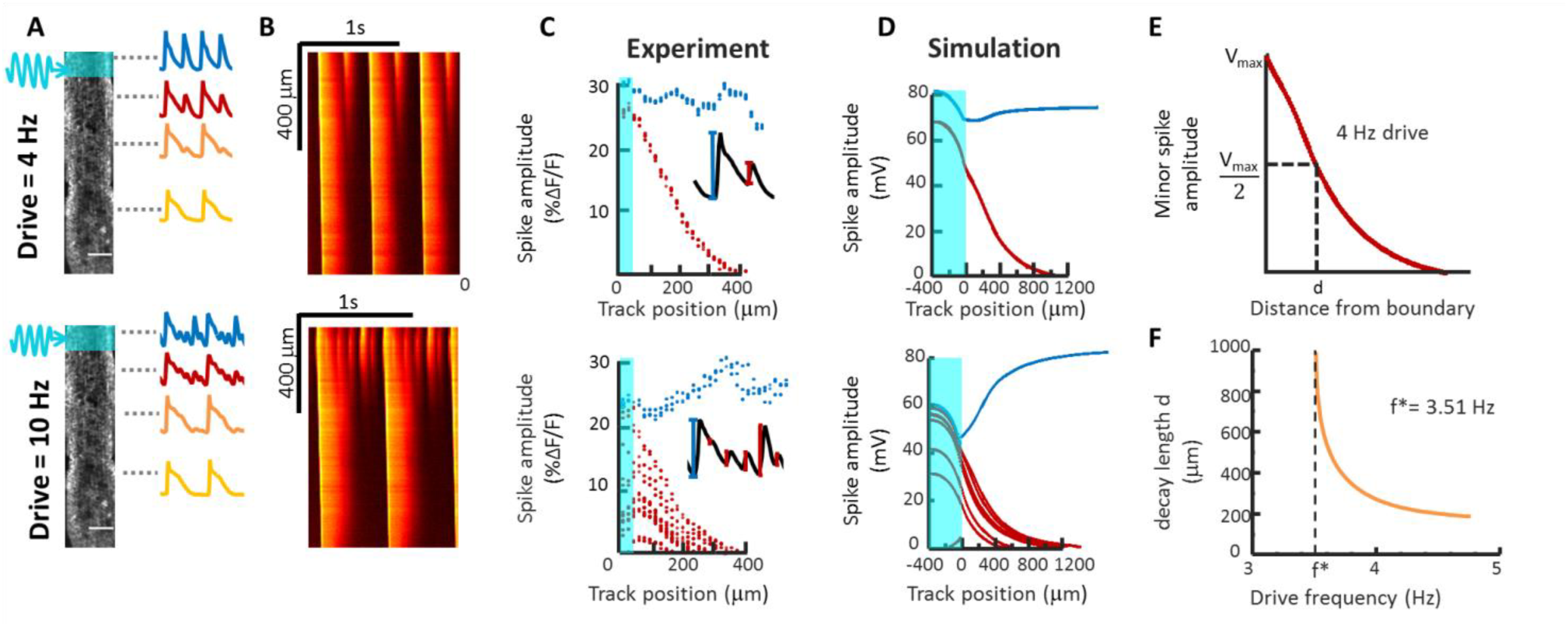
Mapping the transition from near field to far field dynamics. A) 1D tracks were stimulated at one end and action potential waveforms were recorded as a function of distance from the pacing stimulus. B) Kymographs showing electrical dynamics as a function of distance from the pacing stimulus. Some spikes decay in amplitude, while others grow in amplitude. C) Quantification of spike amplitude as a function of distance from the stimulus (shown in cyan). Spikes that reach the far field are shown in blue and spikes that decay are shown in red. D) Numerical simulations under conditions matched to the experiments. E) Definition of the spike decay length, corresponding to 50% loss of amplitude. F) Simulated decay length for near-field spikes as a function of pacing frequency. Below a critical frequency, f*, all spikes reach the far field.

The propagation of spikes in the far-field is governed by the nonlinear dispersion relation, which defines the relationship between wavenumber and frequency (Supplement and Supplementary Fig. 3). The numerically simulated dispersion relation indicated that the far-field dynamics only supported frequencies up to f_max_ = 3.85 Hz. Waves of greater frequency could not propagate into the far field.

At pace frequencies just above the transition to near-field alternans (experimentally observed between 3 and 4 Hz), one might expect period-doubling deviations from a regular spike train to arise slowly. We defined an alternans decay length, d, as the distance over which a near-field alternans beat decayed to 50% of its initial height (Fig. 4e). Simulations near the alternans transition indeed revealed a divergence in *d* near the critical frequency. At pace frequencies far above the critical frequency, the alternans decay length became a small fraction of the far-field action potential length, d ≈ 0.04 *λ* (Fig. 4f), consistent with experimental results (Fig. 4c).

### Second-degree conduction block in iOS-HEK cells

Conduction block and consequent arrhythmias can arise when a region of the heart acts as a partial barrier to conduction.^36^ This effect can arise from spatial variations in either ion channel expression ^37^, or gap junctional coupling.^38, 39^ We thus explored the effect of local defects on spike propagation in iOS-HEK cell tracks. In a serpentine track with sharp turns, we observed that stably propagating far-field waves sometimes failed at the turns (Fig. 5a, **Supplementary Movie 3**). Furthermore, failures occurred in a regular temporal sequence, e.g. Fig. 5a shows a pattern that after the first few beats stabilized into a 3:2 block (3 upstream spikes triggered 2 downstream spikes). The waves developed a curved wavefront and slowed repolarization at the turns, a purely geometrical consequence of the increased electrotonic loading associated with bending a wavefront around a corner. Thus geometrical effects alone are sufficient to cause conduction block, even in a background of homogeneous ion channel levels and gap junction strengths. A related effect has been reported in cultured cardiomyocytes, where a junction of a thin strand of cells to a large island showed unidirectional conduction block due to the inability of the thin strand to drive the large island.^40^

**Figure 5.**
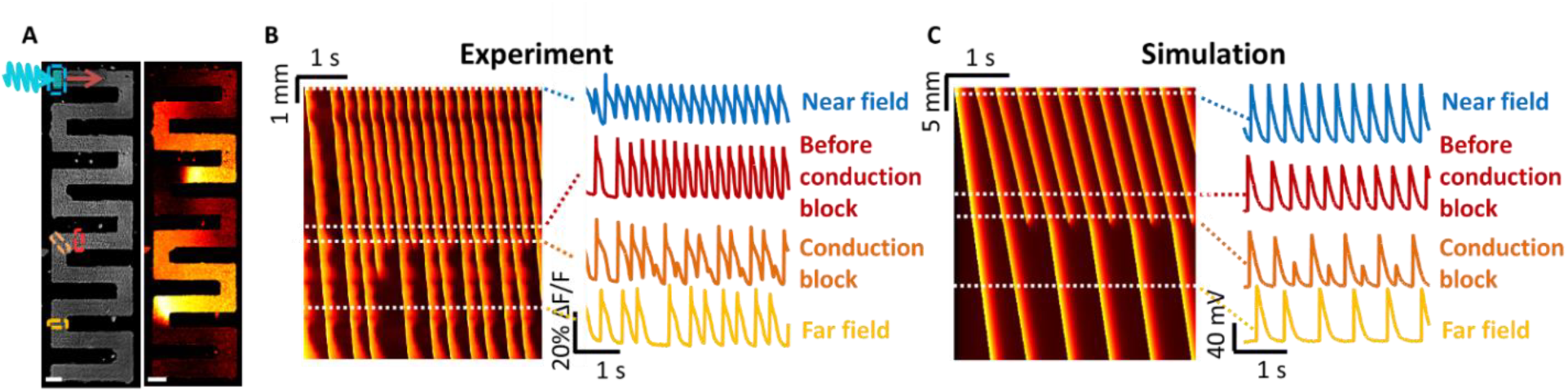
Curvature-induced second-degree conduction block in iOS-HEK cell tracks. A) Left: image of an iOS-HEK cell track showing stimulus region highlighted in cyan and regions of fluorescence monitoring shown with dashed rectangles. Right: single frame from an optical voltage recording showing two action potentials propagating through the track. See **Supplementary Movie 3**. Scale bars 200 μm. B) Left: kymograph showing action potential propagation with a 4 Hz pacing frequency. The vertical axis represents contour coordinate along the track. All spikes conducted into the far-field, but conduction showed second-degree block at a corner where there was a slight defect in the pattern, corresponding to the red and orange rectangles. C) Numerical simulations of second-degree conduction block in a 1D track with a 10 cell zone of 3-fold reduced gap junctional coupling.

In our experiments, the conduction block was attributable to a 2D wavefront curvature effect. To capture this effect in computationally tractable 1D simulations, we simulated linear tracks in which a small region (10 cells) had a reduced gap junctional coupling (*G’_Cxn_ =G_cxn_* / 3). The simulated waves showed a 1:1 conduction block (Fig. 5C), qualitatively similar to that observed experimentally. These experiments and simulations show that local perturbations in the electrotonic coupling, are sufficient to lead to second-degree conduction block.

### Geometry dependent instabilities in human iPSC cardiomyocytes

Finally, we explored whether the geometry-dependent effects observed in iOS-HEK cells also occurred in human iPSC-derived cardiomyocytes (hiPSC-CM). *^41^* Due to the significant commercial interest in using these cells as an *in vitro* model for cardiotoxicity testing ^24, 25, 42-44^, the correspondence (or lack of) between *in vitro* and *in vivo* arrhythmias has some practical importance.

We used microcontact printing to define side-by-side patterns of 0D islands and 1D tracks and then plated hiPSC-CM onto these patterns (Fig. 6b). CheRiff was expressed using a lentiviral vector to allow patterned blue light stimulation^19^, and BeRST1 was used to image changes in membrane potential. We optogenetically paced 0D islands and 1D tracks simultaneously across a range of frequencies. In the 1D tracks, waves propagated with a conduction velocity of 7.2 cm/s and had an AP duration of 360 ms, corresponding to a depolarized action potential length of 2.6 cm.

**Figure 6.**
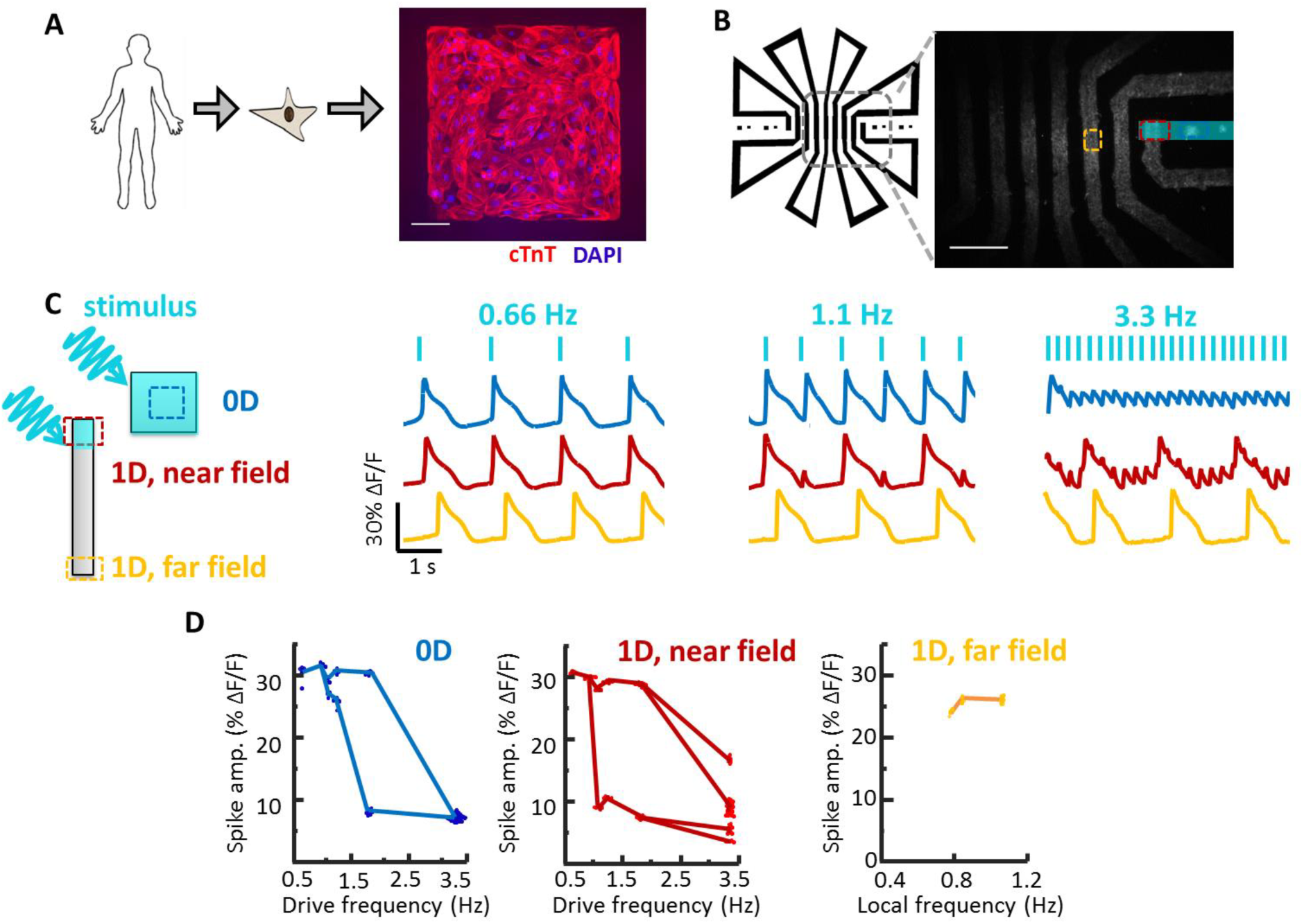
Geometry-dependent arrhythmias in cultured human iPSC-derived cardiomyocytes (hiPSC-CM). A) Image of hiPSC-CM grown on a patterned square island. Scale bar 100 μm. B) Patterned hiPSC-CM cell growth in a set of 0D islands and a long 1D track. The track geometry was designed to interpose long path lengths between returns to the field of view (2 cm between red and yellow regions of interest), to account for the high propagation speed of action potentials in hiPSC-CM compared to iOS-HEK cells. Scale bar 1 mm. C) Simultaneously recorded electrical dynamics in 0D, 1D near field, and 1D far field. At elevated pacing frequencies the cells showed different dynamics in the three geometrical regimes. See **Supplementary Movie 4**. D) Quantification of the action potential height as a function of frequency in the three geometrical regimes.

As with the iOS-HEK cells, we observed that the dynamics depended strongly on the geometry (Fig. 6c, **Supplementary Movie 4**). At high pace frequency (3.3 Hz), the 0D islands showed small regular oscillations, the 1D near-field showed an erratic pattern of large beats with small oscillations superposed, and the 1D far-field only showed the large beats at a sub-harmonic of the pace frequency. As in the iOS-HEK cultures, the alternans beats decayed over a distance much less than the action potential length (Figure 6e; decay length *d* = 535 μm, corresponding to *d* = 0.2 *λ*). Thus the qualitative geometry-dependent behavior of the hiPSC-CM largely mirrored the behavior of the iOS-HEK cells.

The hiPSC-CM cells differed from the iOS-HEK cells in several important regards. First, the hiPSC-CM were spontaneously active, imposing a minimum on the optogenetic pace frequency. Second, the 0D hiPSC-CM islands showed a transition to alternans (Fig. 6c,d) which disappeared at high drive frequencies (3 Hz). Third, the 1D bifurcation to alternans was continuous in the iOS-HEK cells but discontinuous in the hiPSC-CM. In iOS-HEK simulations, these last two differences can be captured simply by tuning the isradipine unbinding rate (Fig. S1b). Simulations of the Noble model also showed clear geometry-dependent differences in the onset of instabilities (Fig. S1c).^45^ Finally, real cardiac tissue can support alternans in far-field traveling waves^46^, while both the iOS-HEK cells and the hiPSC-CM seemed only to support this phenomenon in the near field. Thus there remain important dynamical features of real cardiac tissue which appear not to be captured by either the iOS-HEk cells or the hiPSC-CM.

## Discussion

Gap junction-mediated currents convey the effects of boundaries to all cells in the tissue. The highly simplified iOS-HEK ‘toy’ model revealed that the qualitative dynamics depend in a sensitive way on the overall geometry. Islands of composed of identical cells and differing only in geometry showed vastly different dynamics under identical pacing frequencies, including regular spiking, complex but repeating multi-spike patterns, and irregular spiking. The iOS-HEK system was simple enough to model with biophysically realistic numerical simulations, which confirmed that these observations could be explained by geometry alone. Moreover, the observation of similar geometry-dependent bifurcations in hiPSC-CM cultures suggests that similar principles apply to physiological tissues.

Given the importance of gap junction strength and sample geometry, it is interesting to ask whether one can scale both parameters to preserve the overall dynamics. Such scaling could be useful, for instance, in modeling *in vivo* cardiac dynamics in a cell culture system. We explored this question by systematically varying the gap junction strength in iOS-HEK simulations of a 1-D track. As anticipated from dimensional analysis, the far-field conduction velocity scaled as 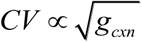 (Fig. S2a). The action potential duration (APD) was largely insensitive to *g_cxn_,* so the action potential length *λ* scaled as 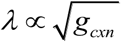 (Fig. S2b). Together these results imply that scaling the size of a system by some factor *k* and the gap junction strength by 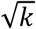 will preserve the overall dynamics.

The transition frequency for alternans, *f^*^*, was largely insensitive to *g_cxn_,* varying by < 2% over a 100-fold change in *g_cxn_* (Fig. S2c), but *f^*^* depended sensitively on the dynamics of the slow recovery variable (Fig. S3). Thus the maximum stable frequency is a parameter that should be largely independent of gap junction strength, and by extension, tissue geometry—provided that the cells are paced via gap junction-mediated conduction rather than by direct pacing.

Can one extrapolate from *in vitro* measurements of small samples with relatively weak gap junction coupling to larger tissues with stronger gap junctional coupling? In iOS-HEK simulations, the alternans decay length, *d*, scaled almost linearly with *λ* (Fig. S2d). These simulations suggest that the action potential length, *λ*, sets a natural length scale for a bioelectric tissue. Moreover, using *λ* as a scaling parameter is advantageous since this parameter can be experimentally measured without knowledge of gap junction conductance or other system parameters.

These results have clear implications for ongoing efforts to predict cardiac stability *in vivo* from *in vitro* systems. Intact myocardium differs in many ways from cultured hiPSC-CM. However, even if one could create hiPSC-CM with fully adult single-cell properties, their behavior in a cultured syncytium would differ dramatically from their behavior in the intact heart. Thus one cannot directly infer *in vivo* behavior from *in vitro* measurements.

The conduction velocity in the atria and ventricles of the human heart *in vivo* is approximately 50 cm/s ^47, 48^, and action potential durations are typically 350 ms, implying an action potential length of *λ* = 18 cm. An adult human heart is approximately *L* = 12 cm long, so L ≈ 0.7 *λ*. Pacemaker-triggered action potentials *in vivo* are thus primarily in the near-field and close far-field regimes. Remarkably, the near-field is the only regime in which we observed arrhythmias in either the iOS-HEK cells or in the hiPSC-CMs, either in experiment or in simulation.

In the cultured hiPSC-CM the action potential length was *λ* = 2.6 cm, so to best match the geometrical regime of the heart, the culture should have size *L* ≈ 0.7 *λ*, or ^~^1.7 cm. Others have reported conduction velocities *in vitro* ranging from 3.5 to 20 cm/s,^21, 33, 49^ corresponding to *λ*= 1.2 to 7 cm (assuming a 350 ms AP width). While cultures of size *L* ≈ 0.7 *λ* are easily accessible on the smaller end of the *λ* range, 5 cm-wide hiPSC-CM cultures are impractical. This challenge can be addressed either by cell patterning to produce serpentine tracks, or by adding gap junction blockers such as 2-APB to diminish the strength of the gap junction coupling and thereby to slow the conduction velocity.

To best mimic activation via the sinoatrial node, cultures of hiPSC-CM should be stimulated locally to launch propagating waves. Whole-field stimuli, e.g. as delivered by field stimulation electrodes or wide-area optogenetic stimulation, will induce synchronous depolarization of all cells, mimicking the 0D case and missing possibly important gap junction-mediated dynamical instabilities. One should ideally perform spatially-resolved voltage measurements to reveal the distinct dynamics in the near- and far-field regimes. While it is not possible to recapitulate the full spatial structure of connectivity of the adult myocardium *in vitro,* one can use the *in vitro* measurements to benchmark dynamical regimes according to biophysical parameters (e.g. *λ* and *L*) which can be measured *in vivo.* We expect that these considerations will be important for ongoing efforts to develop *in vitro* models of cardiac dynamics.

In this work, the use of a simplified synthetic system was crucial to identifying geometry as a determinant of dynamical stability. By comparing experimental results to a biophysically based numerical model, we demonstrated that changing tissue geometry alone could produce dramatically different responses in systems of otherwise identical excitable cells. We anticipate that this synthetic strategy may be useful in clarifying interactions in other complex excitable tissues.^50^

## Materials and methods

### iOS-HEK cell line generation, culture, and patterning

Clonal OS-HEK cell lines were generated as described previously^31^ and maintained in DMEM-10 supplemented with antibiotics to maintain transgene expression. For functional imaging experiments, cells were first incubated for 30 minutes in Tyrode’s solution (containing 125 mM NaCl, 2 mM KCl, 2 mM CaCl_2_, 1 mM MgCl_2_, 10 mM HEPES, 30 mM glucose; pH 7.3, osmolality 305-310 mOsm) supplemented with 10 μg/mL isradipine and 1 μM BeRST1 voltage-sensitive dye.

Patterned cell growth was achieved using previously described methods^28, 32^. Cytophilic patterns of fibronectin were deposited on cytophobic polyacrylamide gels using microcontact printing. Following pattern definition, cells were gently seeded at high density (>500k cells/mL) and allowed to adhere and proliferate throughout the pattern over several days before imaging experiments.

### Wide-field all-optical electrophysiology

All-optical electrophysiology of iOS-HEK and hiPSC-CM cells was performed using an adapted ‘Firefly’ ultrawidefield inverted microscope.^33^ Spatially patterned blue excitation for optogenetic stimulation was achieved using digital micromirror device (DMD). Near-infrared voltage sensors were excited using widefield 635 nm illumination (DILAS 8 W diode laser, M1B-638.3-8C-SS4.3-T3) configured in a near-TIRF configuration Custom LabView software allowed for synchronization of time-modulated signals controlling blue and red light excitation, DMD patterns, and camera acquisitions over experimental runs.

All data were processed and analyzed using custom software (Supplementary Methods). To investigate geometry-dependent dynamical regimes, spatial regions of interest (ROIs) were defined and averaged across pixels to give fluorescence time-traces. A single set of ROIs was defined for a given sample dish and was applied uniformly across movies at different drive frequencies to systematically investigate dynamical responses (Figs. 2B and 6C). Using these spatially resolved measurements, we extract information from multiple replicates of 0D, 1D near-field, and 1D far-field responses all in a single dish.

### Numerical modeling of iOS-HEK cells

In the conductance-based model, the voltage dynamics are governed by the equation

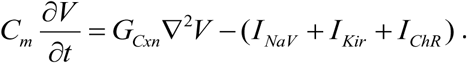

Dynamics of individual currents were modeled according to well-established numerical equations with conductance magnitudes fit to experimental data, and an additional slow gating variable was added to capture activity-dependent sodium channel blockade by isradipine (Supplementary Methods). Numerical simulations were run on isolated cells and on 1D tracks under identical model parameters for comparison to experimental results.

## Acknowledgments

We thank Christopher Werley and Miao-Ping Chien for the hiPSC-CM image in Fig. 6a. This work was supported by the Howard Hughes Medical Institute. HMM was supported by the Department of Defense (DoD) through the National Defense Science & Engineering Graduate Fellowship (NDSEG) program. SD was supported by the NSF Graduate Research Fellowship Program under Grant No 1644760. Y-LH and EM were supported by NIH grant R35GM119855.

## Contributions

HMM and AEC designed the study. HMM conducted and analyzed experiments. HMM and SD developed and simulated the iOS-HEK numerical models. SD and BS conducted dispersion analysis and Noble model simulations. YLH and EWM provided BeRST1 dye reagent. AEC and HMM wrote the manuscript, with input from SD and BS. AEC and BS oversaw the research.

## Supplementary Information

### Supplementary Methods

#### iOS-HEK cell line generation and cell culture

All HEK cells were maintained in Dulbecco’s modified Eagle medium with 10% fetal bovine serum (DMEM-10), penicillin (100 U/mL), and streptomycin (100 μg/mL). OS-HEK cells were generated as described previously.^28^ Briefly, a Na_V_1.5 plasmid (Johns Hopkins University ChemCORE) containing a puromycin selection marker was transfected into HEK cells using TransIT-293 transfection reagent (Mirus Bio). 48 hours after transfection, 2 μg/mL puromycin was introduced into the culture medium for 14 days to select for stably expressing Na_V_1.5 cells. Following this selection, an Optopatch construct containing coding sequences of CheRiff-eGFP and QuasAr2-mOrange (separated by a self-cleaving P2A peptide sequence) was introduced using a lentiviral construct.^29^ GFP-expressing cells were enriched 10 days after infection via fluorescence-activated cell sorting (FACS). Finally, a K_ir_2.1 lentiviral construct containing a blasticidin selection marker was introduced. After 2 days, cell culture media was supplemented with 5 μg/mL blasticidin (to select for K_ir_2.1-expressing cells) and 2 μg/mL puromycin (to ensure continued Na_V_1.5 expression). After 14 days, cells were dispersed as single cells into a 48 well plate, and clonal populations were selected on their ability to spike robustly in response to a blue light stimulus.

Clonal cell lines were maintained in DMEM-10 supplemented with 5 μg/mL blasticidin and 2 μg/mL puromycin in 10 cm tissue culture dishes up until 80% confluence, after which they were trypsinized and cryopreserved at 500,000 cells/vial in 90% DMEM-10, 10% dimethylsulfoxide (DMSO). Cryopreserved vials were then thawed and subcultured in antibiotic-supplemented DMEM-10 in 10 cm tissue culture dishes, and were passaged via trypsinization at 80% confluence at a 1:3 ratio. A given subculture typically maintained robust blue-light induced spiking for approximately seven passages, or 2 weeks, after which cells become less excitable (presumably due to decreased K_ir_2.1 expression over time).

To prepare OS-HEK cells as iOS-HEK cells for functional experiments, cells were incubated in Tyrode’s solution (containing 125 mM NaCl, 2 mM KCl, 2 mM CaCl_2_, 1 mM MgCl_2_, 10 mM HEPES, 30 mM glucose; pH 7.3, osmolality 305-310 mOsm) further supplemented with 10 μg/mL isradipine and 1 μM BeRST1 voltage-sensitive dye for 30 minutes in a mammalian tissue-culture incubator prior to imaging.

#### Cell patterning

Patterned cell growth was achieved using previously described methods^28, 32^. Cytophilic fibronectin patterns were defined on a functionalized cyotophobic polyacrylamide gel using microcontact printing with patterned PDMS stamps. Patterns were first designed *in silico* (Inkscape) and printed onto a Mylar transparency mask (CAD Art Services). Pattern negatives were then transferred onto silicon wafers coated in SU-8 3025 via contact photolithography and subsequent development of unexposed photoresist. PDMS stamps were then cast from the silicon wafer template.

Functionalized polyacrylamide dishes were prepared from MatTek 35 mm-glass bottom dishes. Glass coverslips were chemically activated by plasma cleaning and then incubated for 30 min in a nitrogen-purged glovebox with silane solution (v/v: 0.5% 3-methacryloxypropyltrimethoxysilane, 2% acetic acid, 97.5% anhydrous EtOH). A 40:1 acrylamide:bisacrylamide gel doped with 4.2 mg/mL acryl-NHS (to allow for covalent bonding of fibronectic matrix) was then gelled for 2-3 minutes in ambient air under siliconized coverslips (to ensure smooth gel surfaces). Functionalized acryl-NHS dishes were then sealed in nitrogen and drierite and transferred to −80 C for long-term storage.

To complete dish preparation, patterned PDMS stamps were first coated in fibronectin protein (Yo Proteins no. 663) dissolved in PBS (0.05 mg/mL fibronectin final concentration) in a sterile tissue culture hood. Coating was allowed to settle for 30 minutes, after which excess fibronectin-PBS solution was carefully removed via aspiration. Stamps were further air-dried for 10 minutes to remove excess moisture that could blur pattern transfer. Functionalized dishes were then printed with fibronectin-coated stamps for 1 hour in a tissue-culture incubator, after which stamps were gently removed and dishes re-sterilized for 10 minutes with UV illumination. Cells were deposited by gently pipetting 500 μL DMEM-10 droplet containing the desired cell density (typically between 500,000 and 1M cells/mL). Cells were allowed to adhere for 30 minutes in a tissue culture laminar hood before an additional 2 mL of DMEM-10 was added and cells were transferred to an incubator.

#### Wide-field all-optical electrophysiology

All-optical electrophysiology of iOS-HEK and hiPSC-CM cells was performed using an adapted ‘Firefly’ ultrawidefield inverted microscope.^33^ Spatially patterned blue excitation for optogenetic stimulation was achieved using digital micromirror device (DMD) module with an onboard 460 nm LED (Wintech DLP Lightcrafter 4500). Pixels of linear dimension 7.637 μm were demagnified 2x for optical pattern resolution of approximately 3.8 μm. Action potentials in iOS-HEK cells were triggered with 10 ms pulses at 100 mW/cm^2^; action potentials in hiPSC cardiomyocytes were triggered with 50 ms stimuli. Patterns were defined using custom software (MATLAB). DMD pixels were mapped to sample pixels by calibrating a linear transformation to a test pattern on a fluorescent target.

Near-infrared voltage sensors were excited using widefield 635 nm illumination (DILAS 8 W diode laser, M1B-638.3-8C-SS4.3-T3) configured in a near-TIRF configuration (to reduce background autofluorescence). Robust signals from BeRST1 were obtained at an illumination intensity of 2 W/cm^2^. Near-infrared fluorescence emission was filtered using emission filters (Chroma ET665lp and Semrock quadband 336/510/581/703) and reimaged onto a sCMOS camera (Hamamatsu Orca Flash 4.2). Movies were acquired at 100 Hz over a 5 mm x 5 mm field of view.

Custom LabView software allowed for synchronization of time-modulated signals controlling blue and red light excitation, DMD patterns, and camera acquisitions over experimental runs.

#### Image processing and experimental data analysis

All data were processed and analyzed using custom software (MATLAB). For each pixel, a baseline fluorescence, F, was calculated from the first percentile of the values in the recorded time-trace. The movie was then converted into units of ΔF/F. Pixels that did not contain cells were set to zero (using a criterion that the 99^th^ percentile of a time trace must be over a user-specified threshold). A spatial median filter (5×5 pixel filter) was further applied to account for measurement noise. At the illumination intensities used, BeRST1 photobleaching was negligible.

To investigate geometry-dependent dynamical regimes, spatial regions of interest (ROIs) were defined and averaged across pixels to give fluorescence time-traces. A single set of ROIs was defined for a given sample dish and was applied uniformly across movies at different drive frequencies to systematically investigate dynamical responses (Figs. 2B and 6C). Using these spatially resolved measurements, we extract information from multiple replicates of 0D, 1D near-field, and 1D far-field responses all in a single dish.

We extracted action potential amplitudes for each spike detected in each feature across movies taken at different pacing frequencies. Spike upstrokes were detected via a threshold on the time derivative. Spike amplitudes (in units ΔF/F) were defined as the difference between the maximum value after a given upstroke and the minimum value preceding the upstroke. Between 10 and 300 spikes per feature were collected in a given movie, depending on drive frequency and acquisition time. Beeswarm plots of spike amplitudes at each pacing frequency were used to visualize the dynamical patterns of activity (Figs. 2C, 6E). Only spikes taken from the second half of movies are visualized to avoid early transients which decay as patterns converges to a stable cycle. The drive frequency in 0D and 1D near field measurements was set by the frequency of the blue light-gated CheRiff activation; in 1D far-field measurements, the pacing frequency is defined as the local frequency (i.e., the reciprocal of the interval between successive spikes).

The transition between the near-field and far-field region was mapped using kymographs (Figures 4B and 5B). Kymographs were generated by spatially averaging ΔF/F movies across the short dimension of a linear track. For serpentine tracks (Figure 5), tracks segments were first computationally aligned into a single long track, and then spatially averaged along the short dimension. Alternans transitions were spatially mapped by performing spike detection (as described above) across linear windows of 8 pixels (Fig. 4C).

#### Numerical modeling of iOS-HEK cells

In the conductance-based model, the voltage dynamics are governed by the equation

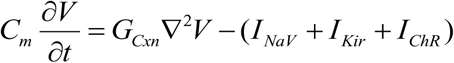

where 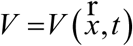 is the voltage in mV. Recall *G_Cxn_* = *g_Cxn_* ×*l^2^*, where *g_Cxn_* is the gap junction conductance between cells, and *l* is the linear dimension of a cell. The ionic currents drive local dynamics, while the diffusion term couples neighboring regions. Units of space are 10^−5^ m (corresponding to linear size of one cell), and time is in ms. Conductances are measured in nS/pF and ionic currents in pA/pF. The currents are:

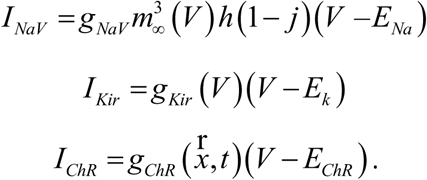

The gating variables for the sodium activation and inactivation gates, *m* and *h* respectively, and for the effect of isradipine, *j,* are dimensionless variables which take values between 0 and 1. The inactivation gate evolves following

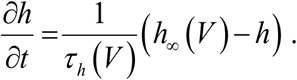

To capture the use-dependent sodium block by isradipine, we introduced a slowly activating and recovering gate, *j*, with behavior governed by

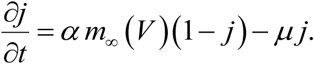

That is, the drug binds sodium channels in its open state with rate *α,* and unbinds with rate *μ.* These kinetic parameters were chosen to fit the experimentally observed dynamical transitions.

Since the time constant of the sodium activation gate, *m*, is orders of magnitude faster than the other gates, the gate was replaced with the asymptotic limit, *m*_∞_ (*V*). This standard approximation allowed for more efficient computations without impacting the simulation results.

The gating variables approach steady-state asymptotic values, which are functions of the voltage.

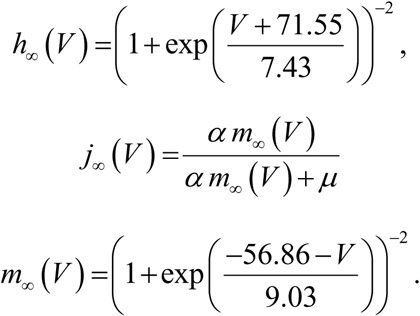

Time constants for each gate tell how quickly the behaviors approach the asymptotic values, and are given by

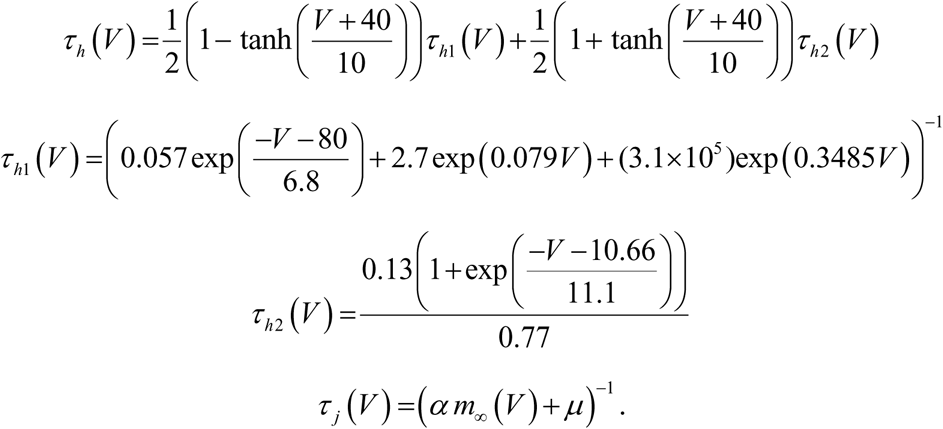

The equations governing the sodium channel activation, *m*, and inactivation, *h*, gates were set following literature values.^34, 35^ Minor modifications were made to the *h* time constant, *τ_h_*(*V*), to smooth the discontinuous function. The channelrhodopsin was modeled as a linear conductance, modulated in space, 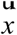, and time, *t,* by the blue light illumination. The repolarization conductance *g_Kir_* was modeled as an instantaneous function of voltage derived from a smoothing spline fit to the action potential waveform at low drive frequency (f = 2 Hz).

Numerical simulations were run on single-cells for the 0D case, and 2 cm 1D tracks with Neumann (no-flux) boundary conditions at each end. The 1D tracks were discretized with a spatial step size, *dx,* equal to 10 *μm*, the length of one OS-HEK cell. With this discretization size, each grid point represented one cell, allowing the simulations to have similar spatial resolution as the experiments.

The Laplacian was modelled with a fourth-order centered finite difference scheme. Boundary conditions were implemented using the standard ghost-point method. A combination IMEX scheme was used for the time evolution, with implicit Crank-Nicholson applied to the diffusion terms and explicit Adams-Bashforth applied to the remaining nonlinear terms (ion currents and gating dynamics). A time step of 0.1*dx* ms (10^−6^ ms in the case of 10 *μm* spatial step) was used, which is well within the accuracies of the numerical scheme and is significantly smaller than any time scales in the model. Single cell, 0D dynamics were modelled by setting *g_cxn_* to zero, and time evolution was performed with the same Adams-Bashforth scheme. All computations were performed using MATLAB.

Initial model parameters were experimentally constrained and fit using a combination of patch clamp measurements and fluorescent voltage recordings. Reversal potentials of *E_Na_* = 75 mV, *E_K_* = −107 mV, and *E_ChR_* = 0 mV were determined according to Nernst equation for the corresponding intracellular and extracellular ionic concentrations. All conductances were expressed as specific conductances per unit capacitance, allowing us to set *C_m_* = 1. The sodium conductance *g_NaV_* was set to 1.5 nS/pF, corresponding to a maximum transient current of 3 nA at −25 mV (*V* – *E_Na_* = –100*mV*) measured in a 20 pF patch clamp recording. The channelrhodopsin conductance was determined to be 15 pS/pF following similar methodology. The inward rectifying potassium current *g_Kir_* was represented as an instantaneous function of voltage, determined by fitting to the experimentally observed fluorescence repolarization waveform at low drive frequency (2 Hz). Because fluorescence traces do not report absolute voltage, we assumed a resting potential of −90 mV and peak voltage of + 30 mV, in accordance with patch clamp recordings. The diffusion coefficient *G_Cxn_* is given by *G_Cxn_* = *g_Cxn_* ×*l^2^*, where *l* = 1 cell. The connexin conductance *g_Cxn_* was estimated according to measurements of the static electronic coupling length^28^ as 40 nS/pF.

The parameters *α* and *μ* were chosen to match experimentally determined dynamical regimes. Setting *α* = 0.5 ms^−1^ and *μ* = 0.015 ms^-1^ gave close agreement with the complex landscape of dynamical transitions observed experimentally (Fig. 3c) as well as simulated voltage traces which closely resembled experimental observations (Fig. 3b). Sensitivity analysis was performed to validate these parameter choices (Supplementary Methods).

Model dynamics were characterized using analogous parameters as for experimental results. Simulations were conducted for both an isolated cell which was directly paced (‘0D’) as well as for linear tracks of 2000 cells (i.e. *L* = 2 cm) in which 40 cells were paced on one end. Simulations were run across a range of stimulus intervals between 500 ms (2 Hz) and 100 ms (10 Hz), sampled at 10 ms intervals. For each numerical experiment, we detected spike upstrokes as upward deflections of the derivative of the simulated

#### Sensitivity analysis of model parameters

Parameter values for *α,μg_Nav_*, and *g_cxn_* were selected to match as closely as possible the experimental results in both the 0D and 1D settings. The model was simulated for 1,500 ms with a given set of parameters under 2 Hz stimuli, and the resulting voltage at times *t_k_*, *V*(*x, t_k_*), was compared to analogous experimental results, *W* (*x,t_k_*), using the ordinary least squares (OLS) objective function

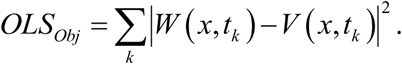

Simulated and experimental voltage values were scaled to be between 0 and 1 to allow for direct comparison. An optimal set of parameters minimizes the objective function; minimization was performed with the MATLAB constrained optimization function fmincon. Realistic parameter constraints were chosen from the patch clamp data.

Sensitivity of model parameters was tested using Latin Hypercube Sampling (LHS)^51^. The LHS results identified *α,μ*, and, *g_NaV_* as the most sensitive parameters, with *g_Cxn_* as the least. That is, small changes in the connexin strength, *g_Cxn_*, had negligible impact on the simulated voltages. Regardless, changing any model parameters over a modest regime maintained the main geometry-dependent differences in stability, giving confidence in the appropriateness of the model.

#### Nonlinear dispersion analysis of far-field dynamics

In the far-field, the waves become spatially periodic structures of constant shape and velocity, both of which are uniquely determined by the frequency. The nonlinear dispersion relation of the far-field equation indicates which wave number (inverse of peak-to-peak wavelength) is selected by each frequency (Supplementary Fig. 3A). The nonlinear dispersion relation is computed by considering the far-field traveling waves as periodic stationary solutions in a moving frame, which allows for the traveling wave and wave number to be solved for as a function of frequency. The full dispersion relation curve is traced out with pseudo-arclength numerical continuation implemented in MATLAB, and using the spatial discretization scheme as in the numerical simulations.

Each frequency has two associated wave numbers, with the bottom branch stable and top branch unstable. Two examples of the spatial structure of the far-field waves are given, and their positions along the dispersion relation marked. The curve does not extend past 3.85 Hz, indicating the far-field equation does not support waves of higher frequencies. Therefore, for drive frequencies above 3.85 Hz, each stimulus will not propagate into the far-field, leading to the appearance of near-field alternans, and aligning with the experimentally and numerically observed behavior.

Changing the drug binding rate, *α*, and connexin strength, *g_cxn_*, have different impacts on the nonlinear dispersion relation (Supplementary Fig. 5B and 5C). Small changes in *α* lead to large shifts in the transition frequency, indicating that faster binding rates will induce near-field alternans at lower drive frequencies. However, large changes to the coupling strength change properties of the traveling waves, such as wave number, wave length, and speed, but do not appreciably change the range of allowed frequencies or transition frequency. Changes to model parameters μ (drug unbinding rate) and *g_Na_* (sodium conductance) are not shown, but have a similar effect as changing *α*.

The 2:1 conduction block observed from local changes in the connexin strength (Fig. 5) is also explained from the nonlinear dispersion relation. As the spike propagates through the small region of lower connexin strength, the wavenumber and frequency are adjusted to values that are no longer stable when the wave reenters the region of higher connexin strength (observe the differences in the curves with *g_cxn_* values of 40 and 14 in Supplementary Fig. 5C). The modification of wavenumber and frequency upon the reentry results in the observed 2:1 conduction block.

#### Noble model of cardiomyocyte dynamics

To demonstrate that the observed geometry effects are not unique to the OS-HEK cell model and are of importance in cardiac dynamics, we performed 0D and 1D simulations in the Noble model. The Noble model is an ionic channel model for Purkinje fibers and the voltage dynamics are governed by sodium, potassium, and background chloride currents (*l_Na_, l_K_*, and *l_cl_*, respectively). Spatial coupling between cells was again incorporated through a Laplacian.

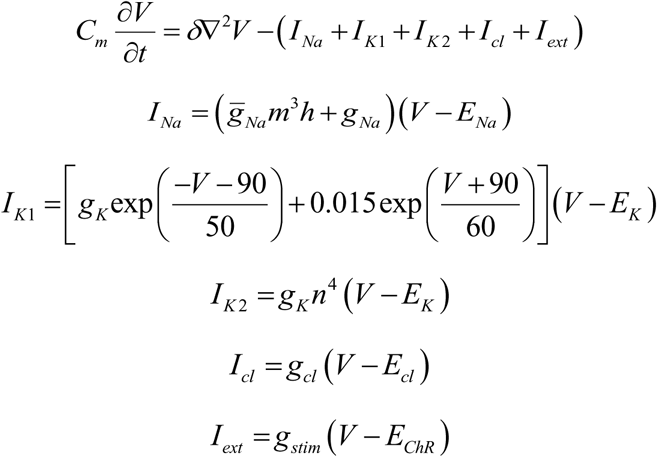

The dynamics of the dimensionless ionic gates for sodium activation, *m,* sodium deactivation, *h,* and potassium, *n,* take the form

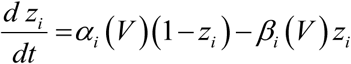

where *z_i_* stands for *m, h*, and *n* and *α_i_* and *β_i_* similarly correspond to opening and closing rates of the *m, h,* and *n* channels. The rates are given by

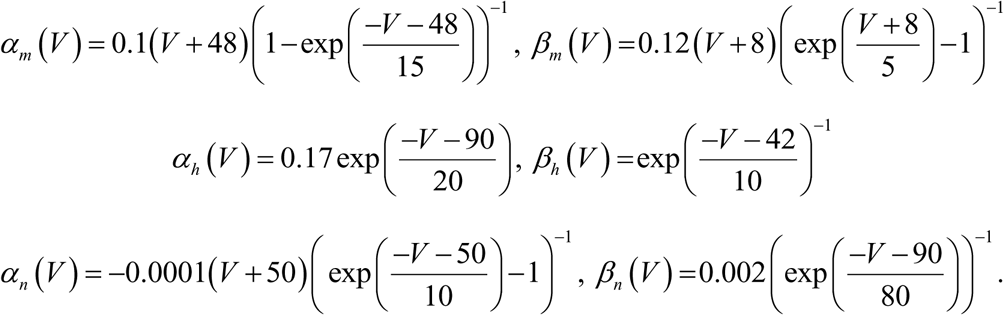

All ionic currents, dynamic gating variables, and parameter values were set from literature. Membrane capacitance, conductances, and Nernst potentials are given in Table 1.^45, 52^ The form of the applied current, *I_ext_*, was set to mimic the channelrhodopsin stimulus. The Noble model was chosen for its ability to reproduce realistic behaviors observed in cardiac cells and the relatively low number of ionic currents allowed for efficient simulation.

**Table 1:**
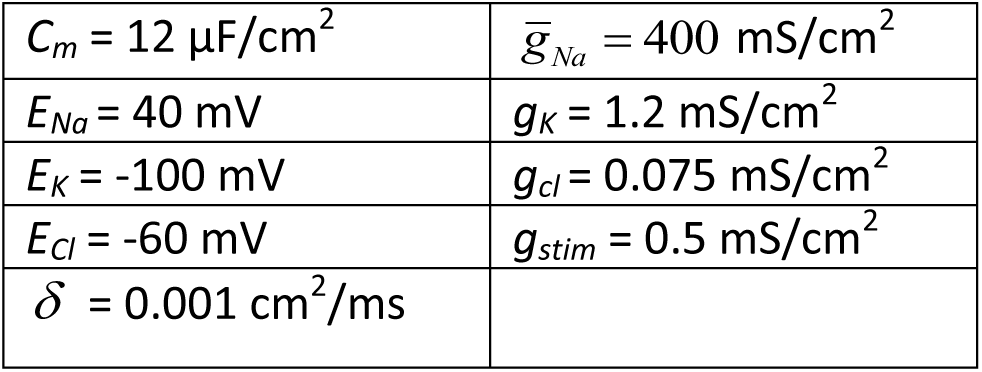
Values of membrane conductance, conductances, and Nerst potentials used in simulations of the Noble model.

Numerical simulations in 0D and 1D geometries used the same centered finite-difference Laplacian and IMEX time evolution schemes described above. Simulations were run on 1D tracks of up to 6 cm in length with spatial grid spacing set to 0.1 mm and time step of 0.01 ms. Due to the higher cell coupling and conduction values, the stimulus duration was increased to 50 ms. Simulations were run for at least two forcing cycles before recording results to remove any transient effects caused by the self-spiking behavior of the Noble model. Model output was analyzed in a similar fashion to the OS-HEK model results.

The Noble model simulations also exhibit geometry-dependent changes in stability (Supplementary Fig. 1C). The 1D near-field results show a dramatic transition first to alternans at 4.5 Hz, and then to chaotic-like dynamics at 8 Hz. Similar transitions are not observed in the 0D or far-field results, which instead display monotonically decreasing spike amplitude for higher frequencies. As before, the model selects a narrow range of wave frequencies which propagate into the far-field.

## Supplementary Figures

**Figure S1.**
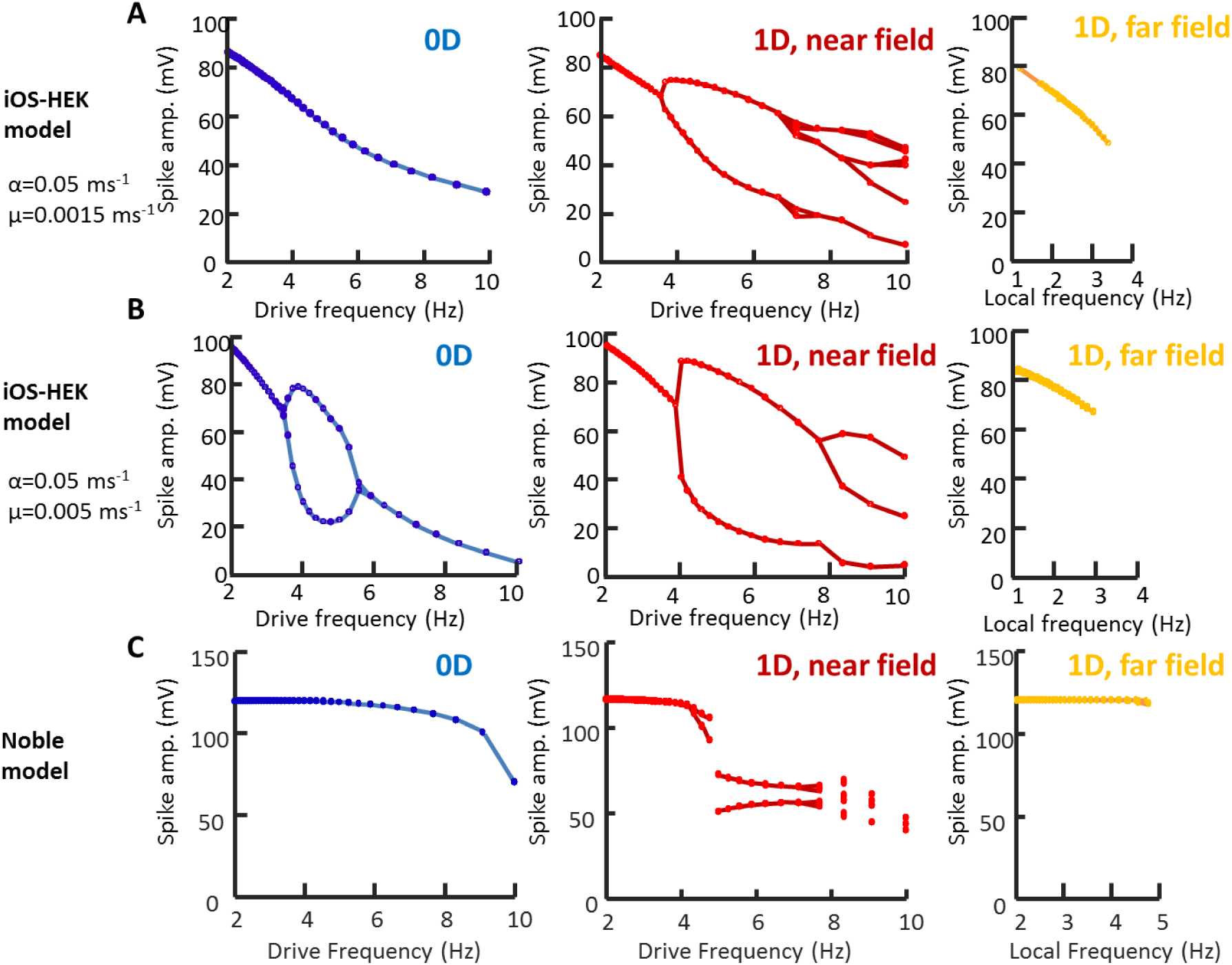
Geometry dependent instabilities in different numerical models of excitable cells. A) Hodgkin Huxley model of iOS-HEK cells (same as Fig. 3c, repeated here for comparison to other models). B) Same as (A) with artificially accelerated israpidine unbinding kinetics (μ = 0.005 ms^−1^). 0D features show alternans at intermediate drive frequencies. The 1D near field shows a discontinuous alternans transition. C) Simulation of the cardiac Noble model ^45^, which also shows geometry-dependent changes in dynamics.

**Figure S2.**
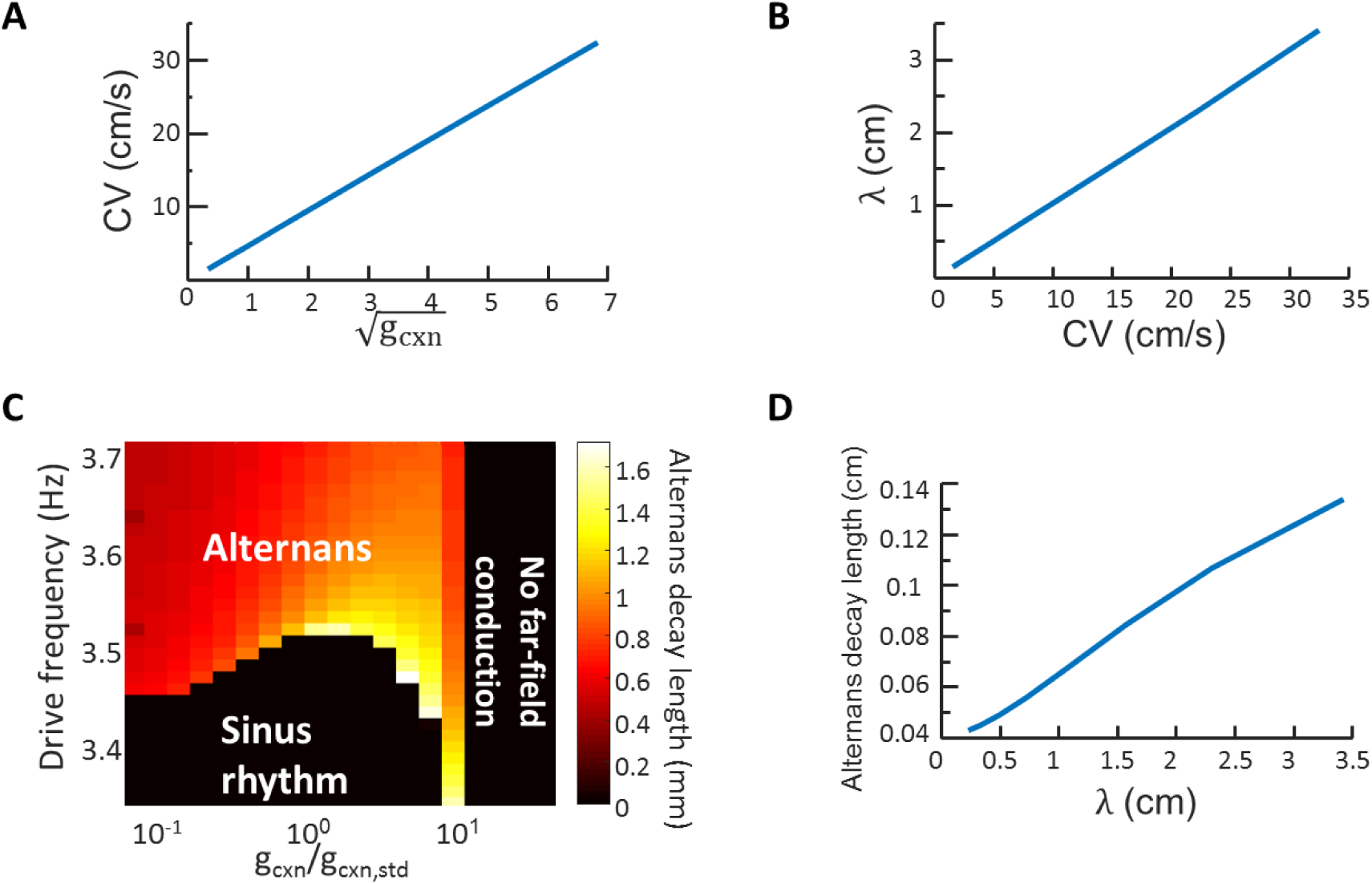
Effect of gap junction coupling on dynamics and instabilities in Hodgkin-Huxley simulations of iOS-HEK cells. A) Linear scaling of conduction velocity with √ *g_cxn_*. B) Linear scaling of depolarized pulse length, *λ*, with conduction velocity. C) Frequency of near-field alternans onset as a function of gap junction coupling strength. The onset frequency varies by < 2% over a 100-fold variation in *g_cxn_.* Pseudocolor shows the spatial extent of alternans (measured as the distance over which alternans beats decay to half of their value in the stimulus region). D) Relation of the alternans decay length to propagating pulse wavelength at a pacing frequency of 5 Hz, far above the onset of alternans.

**Figure S3.**
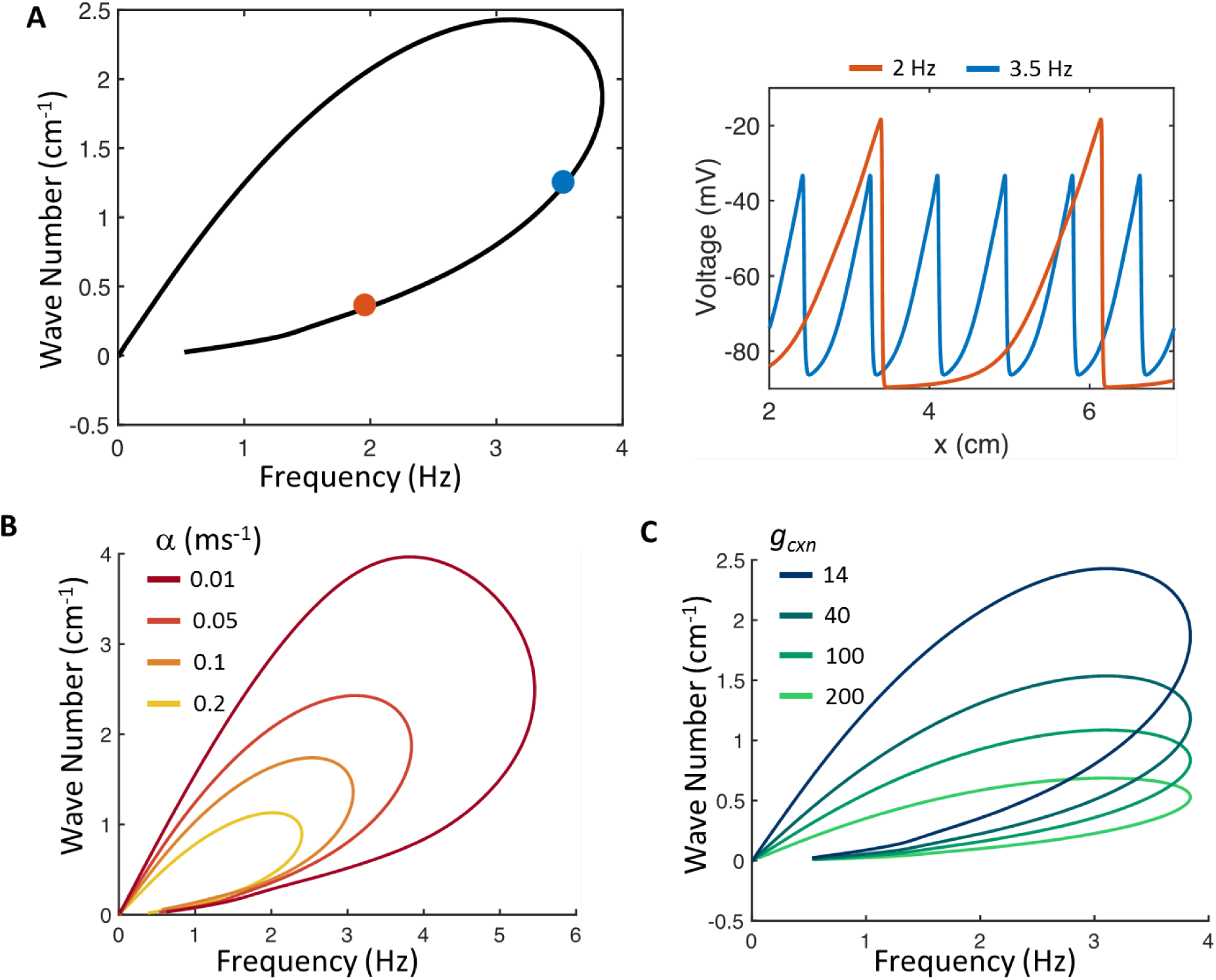
Dispersion relation for iOS-HEK cell model. Nonlinear dispersion relations show which frequencies are stable in far-field and give wave numbers (inverse of peak-to-peak wavelength) selected by each frequency. In all figures, bottom branches of nonlinear dispersion relations are stable and the top branches are unstable. A) Left: nonlinear dispersion relation under conditions corresponding to experimental data. Right: spatial structure of voltage waves in the far-field with frequencies 2 and 3.5 Hz. Position on nonlinear dispersion relation indicated by correspondingly colored dots. B) Effect of varying the isradipine binding rate, *α*, on the nonlinear dispersion relation. Increasing *α* leads to a smaller range of frequencies that propagate to the far field. C) Changes in the connexin strength, *g_cxn_,* affect the wavelength but have little effect on the maximum frequency prior to alternans onset. Thus the maximum frequency prior to alternans onset is a parameter that is largely independent of gap junction strength, and by extension, tissue geometry.

## Supplementary Movies

**Movie S1. Electrical waves propagating through patterned iOS-HEK cells**. Optical recordings of optogenetically triggered action potentials propagating through confluent monolayers of patterned electrically excitable iOS-HEK cells. See Fig. 1 for details.

**Movie S2. Geometry dependent action potential dynamics in iOS-HEK cells**. The square islands on the right produced spikes in response to each stimulus. The 1D tracks on the left produced an alternans pattern in the near-field and conducting waves only on alternating stimuli.

**Movie S3. Second degree conduction block near sharp turns in an iOS-HEK serpentine track**. The first four action potentials conducted into the far-field, but the fifth failed to conduct at a defect in the middle of the track. Thereafter the waves showed a 3:2 conduction block at the defect in the track.

**Movie S4. Geometry dependent action potential dynamics in hiPSC-CM**. The square islands on the right produced spikes in response to each stimulus. The 1D tracks on the left produced an alternans pattern in the near-field and conducting waves only on alternating stimuli.

